# *Leishmania* targets the macrophage epigenome and dampens the NF-κB/NLRP3-mediated inflammatory response

**DOI:** 10.1101/649632

**Authors:** Hervé Lecoeur, Eric Prina, Thibault Rosazza, Kossiwa Kokou, Paya N’Diaye, Nathalie Aulner, Hugo Varet, Giovanni Bussotti, Yue Xing, Robert Weil, Guangxun Meng, Gerald F. Späth

**Affiliations:** Institut Pasteur, INSERM U1201, Unité de Parasitologie moléculaire et Signalisation, Département des Parasites et Insectes vecteurs, 25 Rue du Dr Roux, 75015 Paris, France; Key Laboratory of Molecular Virology and Immunology, Institut Pasteur of Shanghai, Chinese Academy of Sciences, Shanghai, China; Institut Pasteur International Mixed Unit ‘Inflammation and *Leishmania* infection’; Institut Pasteur, Photonic BioImaging (UTechs PBI), 28 Rue du Dr Roux, 75015 Paris, France; Institut Pasteur, Hub Bioinformatique et biostatistique, 28 Rue du Dr Roux, 75015 Paris, France; Institut Pasteur, Laboratoire de Signalisation et Pathogenèse, 25 Rue du Dr Roux, 75015 Paris, France

**Keywords:** *Leishmania amazonensis*, amastigotes, NF-κB, NLRP3 inflammasome, histone H3 acetylation

## Abstract

Aberrant macrophage activation during intracellular infection generates important immunopathologies that can cause severe human morbidity. A better understanding of microbial immune subversion strategies and macrophage phenotypic and functional responses is a prerequisite for the design of novel, host-directed intervention strategies. Here, we uncover a fine-tuned transcriptional response induced in primary macrophages infected by the human parasite *Leishmania amazonensis* that prevents NF-κB and NLRP3 inflammasome activation. This unusual subversion is characterized by respectively suppression and induction of activating and de-activating components of the NF-κB and NLRP3 pathways. This dichotomic modulation was associated with histone H3 hypoacetylation at promoters of NF-κB-related, pro-inflammatory genes. Our results reveal a novel *Leishmania* immune subversion strategy targeting host cell epigenetic regulation to modulate the macrophage phenotype. Modulation of the macrophage epigenetic landscape establishes conditions beneficial for intracellular parasite survival, and opens interesting new venues for host-directed, anti-microbial drug discovery.

## Introduction

Macrophages are phagocytic cells of the reticulo-endothelial system that carry out a plethora of functions, ranging from maintenance of tissue-specific homeostasis / repair to anti-microbial activity, or involvement in appropriate immune responses in cancer or infection (Ginhoux and Jung, 2014; Gordon and Taylor, 2005). This functional diversity is matched by an equally diverse phenotypic landscape, with macrophages acquiring distinct and unique morphological and functional properties in response to tissue-specific determinants or interactions with tumor cells and microbes (Biswas and Mantovani, 2010; Gordon and Martinez, 2010). While this plasticity adapts the macrophage response to very diverse physiological and microbial insults, aberrant polarization has been linked to various pathologies, in particular intracellular infection (Murray et al., 2014). Many viral, bacterial and eukaryotic pathogens exploit macrophages as host cells and have co-evolved strategies to modulate the macrophage phenotype promoting their own survival (Benoit et al., 2008; Gazzinelli et al., 2014; Herbein and Varin, 2010; Herbert et al., 2004; Noel et al., 2004; Pathak et al., 2007; Pearce and MacDonald, 2002; Raes et al., 2007). Intracellular pathogens thus are a major threat to human health, but also represent interesting tools to probe macrophage plasticity and understand its role in the pathophysiology of infection. This is well exemplified by the intracellular parasite *Leishmania* that has been used as model pathogen to gain insight into innate and acquired immune response pathways that govern resistance or susceptibility to microbial infection (Bellamy, 1999; Canonne-Hergaux et al., 1999; Sacks and Noben-Trauth, 2002; Scott and Farrell, 1981, 1982).

Protist pathogens of genus *Leishmania* cause severe immunopathologies in humans and animals termed leishmaniases (Alvar et al., 2012). Following transmission of the insect-stage promastigote form of the parasite to vertebrate hosts by blood-feeding infected sand flies, the parasite develops into the amastigote form inside the fully acidified phagolysosomes of host macrophages (Antoine et al., 1998; Zilberstein and Shapira, 1994). Intracellular amastigotes manipulate host cell signaling, immune functions, and metabolism to establish permissive conditions for long-term chronic infection (Arango Duque and Descoteaux, 2015; Giraud et al., 2012; Lecoeur et al., 2010; Lecoeur et al., 2013; McConville et al., 2015; Olivier et al., 2005; Osorio y Fortea et al., 2009; Osorio y Fortea et al., 2007). In recent years, the impact of intracellular *Leishmania* infection on the NF-κB-mediated and inflammasome-dependent pro-inflammatory response has attracted important attention. NF-κB denotes an eukaryotic protein family of related transcription factors that form various homo- and heterodimeric complexes involved in regulating promoter activities during inflammation, including the pro-inflammatory cytokines interleukin-1β (IL-1β), interleukin-18 (IL-18) and the NLRP3 inflammasome (He et al., 2016). The inflammasome represents a series of related multi-protein complexes defined by specific Nod-like receptor proteins (NLRPs) that lead to caspase 1-dependent maturation of pro-IL-1β and pro-IL-18 (Franchi et al., 2012), thus placing this pathway downstream of NF-κB (Jo et al., 2016). Despite the plethora of studies investigating *Leishmania* anti- and pro-inflammatory activities, only little is known on the mechanisms that modulate NF-κB-regulated and inflammasome-dependent gene expression in *Leishmania* infected macrophages, and how the plasticity of the macrophage response is exploited by this pathogen to favor its intracellular survival.

Here, by using a physiologically relevant *ex vivo* experimental system of murine primary macrophages infected with lesion-derived *L. amazonensis* amastigotes, we provide first insight into a novel immune subversion mechanism based on histone H3 post-translational modifications, which may have escaped previous investigations that were largely conducted using *Leishmania* promastigotes (Dey et al., 2018; Gurung et al., 2015; Lima-Junior et al., 2013). Indeed, we show that only amastigotes cause transcriptional inhibition of the host cell pro-inflammatory responses via dichotomic regulation of activators and inhibitors of the NF-κB - NLRP3 axis. Our results reveal a novel *Leishmania* immune subversion strategy targeting host cell epigenetic regulation to modulate macrophage phenotypic plasticity beneficial for intracellular parasite survival, thus opening interesting new venues for host-directed, anti-microbial drug discovery.

## Results

### *L. amazonensis* amastigotes prevent inflammasome priming and activation

The macrophage inflammasome response to *L. amazonensis* (*L. am*) infection has been previously studied using insect-stage promastigotes derived from *in vitro* culture (Lima-Junior et al., 2013). Here we assess the impact of disease-relevant, bona fide *L. am*. amastigotes on inflammasome priming and activation in primary, bone marrow-derived, murine macrophages. Unlike LPS/ATP-treated controls, BMDM infection with lesion-derived *L. am* amastigotes did not lead to inflammasome activation as judged by the absence of secretion of mature IL-1β and active caspase 1 p20 (Figure 1A) and ASC oligomerization (Figure 1B). On the contrary, infection caused a significant reduction of all major inflammasome receptor proteins, including NLRP3, NLRC4, AIM-2 and RIG-I (Figure 1C). This was consistent with a similar reduction in transcript abundance in response to live but not heat-killed parasites or inert latex beads (Figure 1D). Down-regulation was specific to amastigotes as judged by normal abundance of *il1β, il18* and *nlrp3* transcripts 24h after infection with purified metacyclic promastigotes, which resulted in reduction of abundance of these transcripts only at day 3 PI after parasite differentiation into amastigotes (Figure S1). Decreased levels of *il1β* and *il18* transcripts correlated with increased amastigote burden during long term cultures (30 days of infection, Figure 1E and 1F, Figure S1C). Significantly, long-term infection had no effect on macrophage phagocytic activity, surface marker expression (Figure S1D), or transcript expression of various inflammasome-independent cytokines, including *il1 α* and *tnf* (Figure S1E, S1F), while a persistent down modulation of transcripts for additional inflammasome components was induced (Figure S1G).

**Figure 1:**
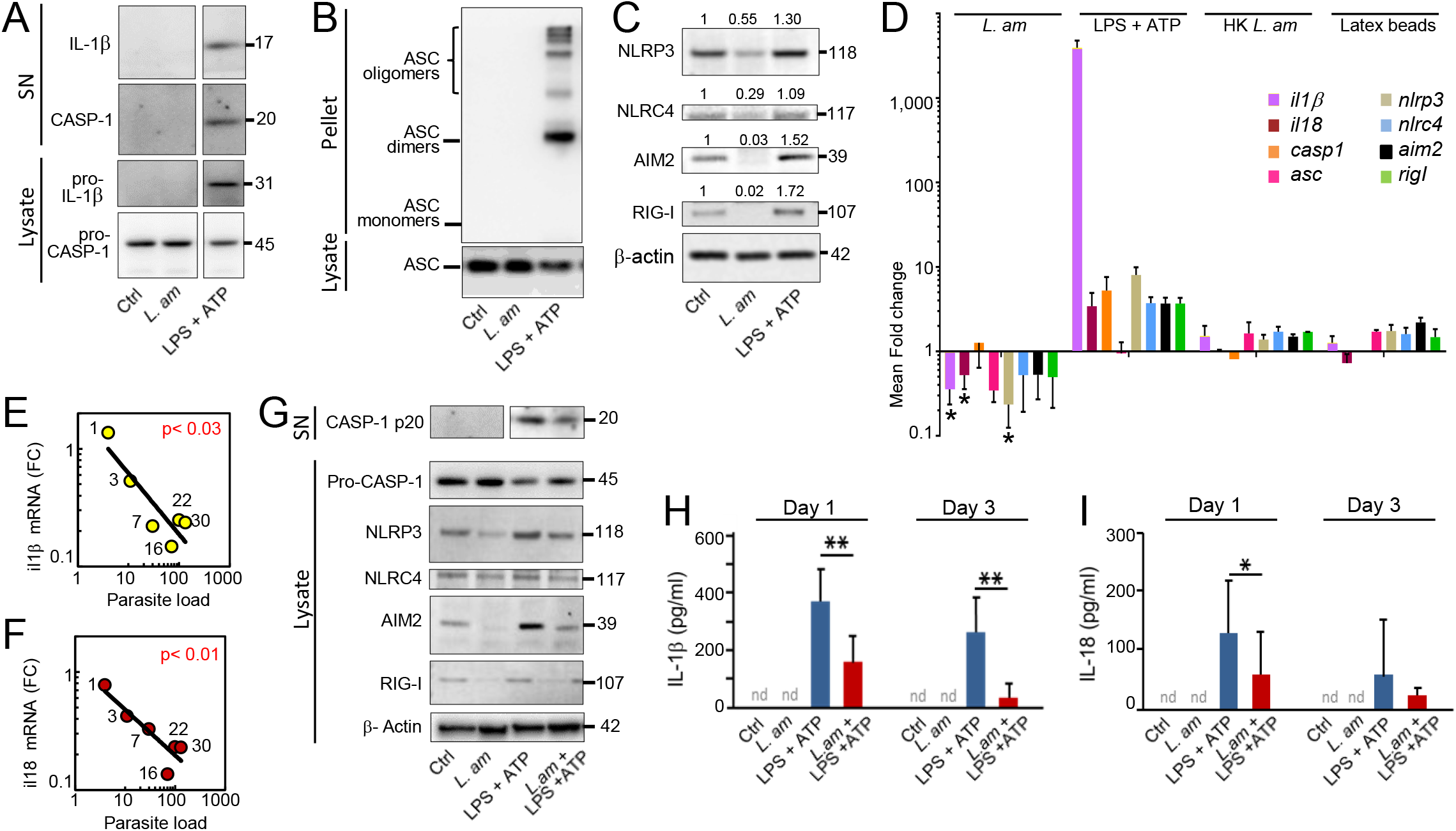
Prevention of inflammasome priming and activation in *L. am*-infected macrophages. BMDMs were infected with amastigotes at a MOI of 4:1. (A) Western blot analysis of mature IL-1β and caspase-1 p20 in culture supernatants (SN), and pro-caspase-1 and pro-IL-1β in macrophage lysates (day 3 PI). (B) Western blot analysis of ASC in cell lysates and pellets (day 3 PI). (C) Western blot analysis of NLRP3, NLRC4, AIM-2 and RIG-I inflammasomes in BMDM lysates (day 3 PI). Beta-actin was used as a control protein. (D) Transcriptional modulation of inflammasome components as assessed by RT-qPCR. BMDMs were loaded with live (*L. am*) or heat-killed (HK *L. am*) amastigotes, or latex beads and cultured for 3 days. As positive control, NLRP3 activation was induced after LPS and ATP stimulation. The expression fold changes (FC) are indicated using uninfected and unstimulated macrophages as a calibrator. Histograms display mean FC values ± SEM for *L. am*-infected and LPS + ATP-treated uninfected BMDMs (n = 4-13 independent experiments) and for HK *L. am* and latex beads-loaded BMDMs (technical duplicates, one representative experiment is shown). (E, F) Biparametric dot plots displaying transcript modulation (fold changes, FC) of IL-1β (E) and IL-18 (F) and parasite load (mean number of amastigotes per macrophage) at different time points PI (indicated by the numeric label of the data points). The F-test p-value is indicated for the corresponding linear regression. (G) Western Blot analyses of inflammasome components at day 3 PI in culture supernatants (SN) and lysates from LPS-stimulated and non-stimulated samples. (H, I) Detection by ELISA of secreted IL-1β (H) and IL-18 (I) at day 1 and 3 PI. Blue and red bars represent cytokine levels for uninfected and infected BMDMs, respectively (mean +/− SEM, n = 3-6). nd, not detected; *, p < 0.06; **, p < 0.05.

The subversion of anti-inflammatory functions of macrophages by *L. am* amastigotes was also observed in response to LPS and ATP treatment, which are used to induce NLRP3 priming and activation. *L. am* infection interfered with NLRP3 inflammasome activation on multiple levels by (i) reduction of physiologically active caspase 1 p20 in the supernatant and the increased expression of inactive pro-caspase 1 in the cytoplasm (Figure 1G), (ii) decrease in abundance of NLRP3, RIG-I, and AIM2 inflammasome (Figure 1G), and (iii) reduction of IL-1β and IL-18 secretion (Figure 1H and 1I), which was independent of the duration and concentration of ATP stimulation (data not shown). Again, in agreement with our observations in unstimulated macrophages, subversion of inflammasome priming and activation in LPS/ATP stimulated macrophages was amastigote-specific, since metacyclic promastigotes did not interfere with the secretion nor the transcriptional modulation of IL-1β nor IL-18 until their differentiation into amastigotes (day 3 PI) (Figure S2).

Interestingly, a different pattern was observed for TNF and IL-1α, two pro-inflammatory cytokines whose production and secretion are independent of inflammasome activation: While *L. am* amastigote infection did not modify IL-1α secretion nor transcript stability (Figure S3A and S3C), it synergized with LPS/ATP for the production of pro-inflammatory TNF (Figure S3B), apparently as a result of an increased stability of *tnf* mRNA (Figure S3D). In conclusion, our data demonstrate that *L. am* amastigotes specifically inhibit the basal and LPS/ATP-induced expression level of inflammasome components and their target cytokines by an active, parasite-driven and stage-specific mechanism.

### Pleiotropic inhibition of NF-κB activation by *L. am* amastigotes

We next investigated the impact of *L. am* amastigote infection on the activity of NF-κB p65 (RelA), a master regulator of the inflammatory response that has been shown crucial for the priming step of NLRP3 activation and IL-1β and IL-18 expression (Jo et al., 2016). The analysis of the NF-κB pathway in uninfected and *L. am*- infected BMDMs revealed inhibition of this pathway by the intracellular parasites at multiple levels. First, using a high content quantitative imaging approach we observed a highly significant reduction of the nuclear localization of RelA in response to infection at 30 and 120 min of LPS treatment (Figure 2A and 2B). Second, we revealed a two-fold decrease in total RelA protein expression in infected cells regardless of LPS treatment (Figure 2C), which partially explained its reduced nuclear presence. Third, infection interfered with the phosphorylation of the inhibitor of NF-κB translocation (IκBα) in response to LPS stimulation as revealed in samples treated with the proteasome inhibitor MG132 that prevents proteasomal degradation of phosphorylated IκB (P-IκBα) (Figure 2D). Reduced IκBα phosphorylation was correlated with a strong reduction of the IκB protein kinase (IKKβ) expression in *L*. am-infected BMDMs (Figure 2D). As judged by RT-qPCR, the decreased protein level is the consequence of the reduced expression of *rela, iκbα* and *ikkβ* transcripts evidenced in absence (Figure 2E) or presence (Figure 2F) of a short pulse (30 min) of LPS. Strong inhibition was also observed for other NF-κB family members (*relb* (p50), *nf-κB1* and *nf-κb2*), the NF-κB inhibitors *iκbβ* and *iκbε*, and the NF-κB activators *ikkα* and *ikkγ*, and persisted during long-term infection (Figure S4).

**Figure 2:**
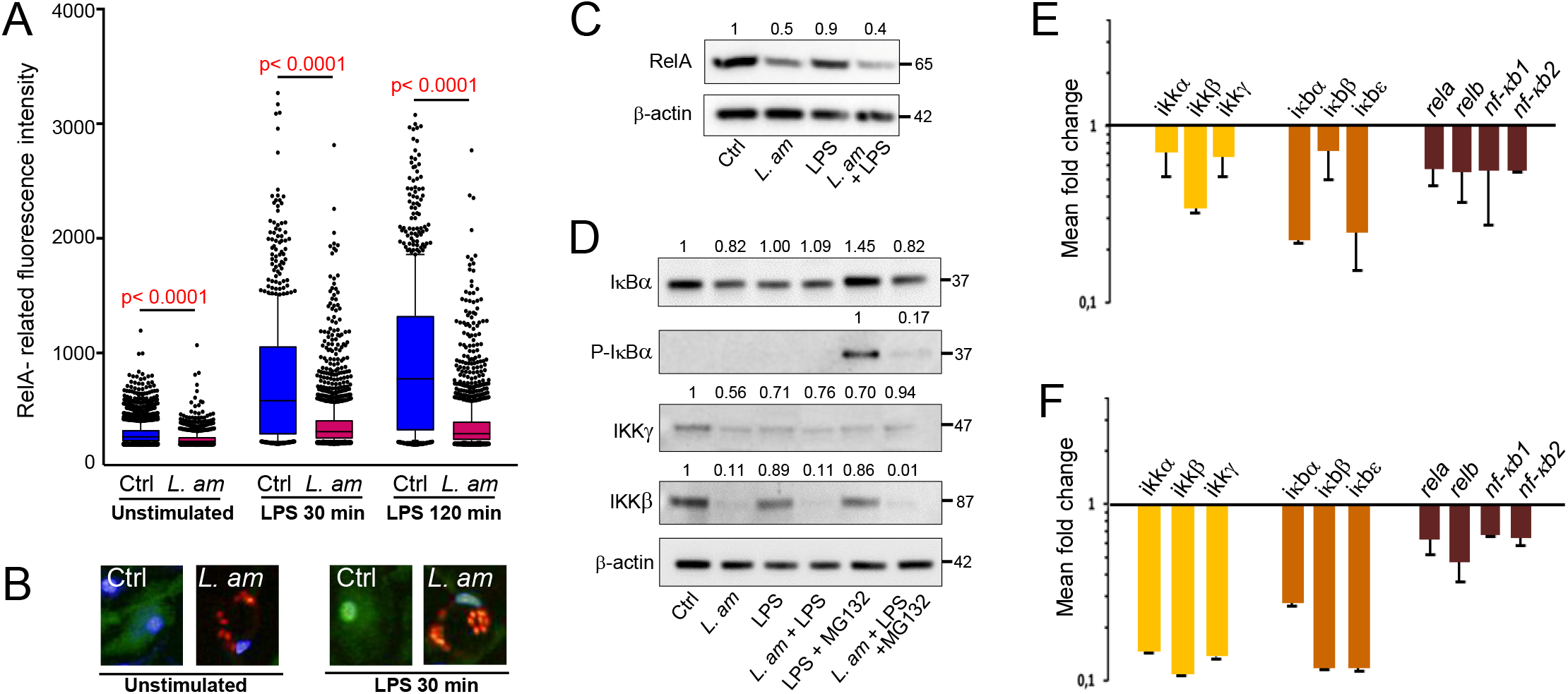
Subversion of the NF-κB pathway in *L. am*-infected BMDMs. BMDMs were infected by lesional *L. am* amastigotes. At 3 days PI, uninfected (Ctrl) and infected (*L. am*) BMDMs were stimulated or not with LPS for 30 or 120 min. (A) Quantitative evaluation of nuclear translocation of RelA (p65) by high content imaging. RelA was evidenced using an anti-RelA polyclonal antibody and an Alexa Fluor™ 488 conjugate (green staining). Cell nuclei were stained with Hoechst 33342 and parasites were visualized by mCherry fluorescence (red staining). Image acquisition and analysis were performed using the OPERA QEHS and the Columbus image storage and analysis system, respectively. The mean fluorescence intensity of RelA-associated staining in BMDM nuclei after 30 and 120 min of LPS stimulation is shown (blue and red box-and-whisker plots, respectively) (800 < analyzed nuclei < 2000) (A). Representative images are shown (B). Note the moderate green (RelA-related) staining in nuclei from infected versus uninfected BMDMs. (C, D) Relative quantitation by Western blot analysis of selected key actors of the NF-κB pathway in total protein extracts. At 3 days PI, BMDM cultures were stimulated sequentially with LPS (4 hours) and ATP (2 hours). The following proteins were analyzed: RelA (C), IκBα and its phosphorylated form (P-IκBα), IKKγ and IKKβ (D). MG132 was added to prevent proteasomal degradation allowing for the revelation of phosphorylated IκBα. (E, F) Transcript modulations of NF-κB members in *L. am*-infected BMDMs. RT-qPCR was performed in samples without LPS (E) (n = 3 experiments) or after 30 min of LPS stimulation (F) (n = 1 representative experiment). Results are shown as fold change values obtained between infected and uninfected BMDMs (calibrator) for members of the signalosome (*ikkα, ikkβ* and *ikkγ*, yellow bars), kinases (*iκbα, iκbβ, iκbε*, orange bars) and NF-κB family members (*rela, relb, nfκb1, nfκb2*, brown bars).

### Transcript profiling reveals dichotomic inhibition of the NF-κB-NLRP3 axis by *L. am* amastigotes

Given the pleiotropic effect of *L. am* amastigotes on the expression of various components of the NF-κB pathway, we extended our transcript analysis using RT-qPCR to investigate the impact of *L. am* infection on the expression of 97components of the NF-κB/NLRP3 axis (Figure 3). We revealed a surprising dichotomic transcriptional pattern: On one hand, infection induced a global down-modulation of transcripts for positive regulators of the NF-κB pathway, including the pro-inflammatory surface receptors *il18r1* (Log2FC =-1.83), *tnfrsf1A* (Log2FC=-1.04), *tlr4* (Log2FC=-0.64), the *myd88* adaptor (Log2FC=-0.57) and kinases such as *irak4* (Log2FC=-0.77) and *mapk14* (Log2FC=-0.98). On the other hand, infection up regulated transcripts of anti-inflammatory molecules and known inhibitors of NF-κB activation such as *tollip* (Log2FC=+0.95) or *otud7b* (Log2FC=+2.08) (Figure 3A and Figure S5). A similar regulatory dichotomy was observed for a series of known NLRP3 activators (*p2rx7;* Log2FC= −1.63) and inhibitors (*tnfaip3*; Log2FC=+1.84) (Figure 3B).

**Figure 3:**
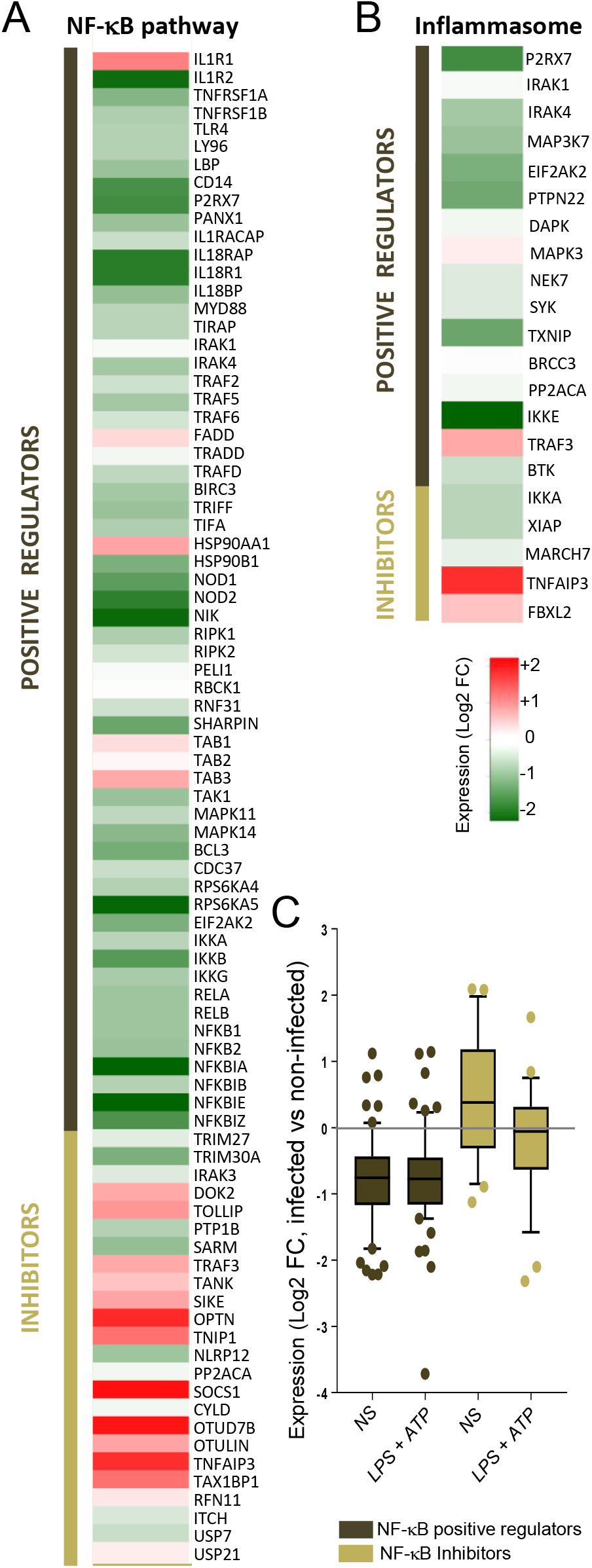
Transcriptional regulatory dichotomy of the NF-κB signaling pathway in *L. am-* infected macrophages. After 3 days PI, total RNA from uninfected and *L. am*-infected BMDMs was isolated and reverse transcribed. cDNA was subjected to RT-qPCR analysis targeting components of the NF-κB and NLRP3 pathways. Log2 values of fold changes in transcript abundance are represented as heat-maps (infected versus uninfected BMDMs) for activators and inhibitors of the NF-κB (A) and inflammasome (B) pathways (mean +/− SD, 3 < n < 8 independent experiments). (C) Comparison of global transcriptional modulation induced by *L. am* parasites in unstimulated (NS) or LPS/ATP stimulated BMDMs for activators (dark brown) and inhibitors (light brown) of the NF-κB pathway. Box- and Whisker plots represent the mean and the 10-90 percentiles of the Log2 fold change values.

A slightly different pattern was observed in infected cells stimulated by LPS/ATP: while the down-modulation of NF-κB positive regulators remained robust, no significant effect on inhibitory components was observed (Figure 3C and Figure S5), which may explain the dampening rather than abrogation of inflammasome activation we observed in LPS/ATP-stimulated, infected BMDMs. Together these data uncover a surprisingly fine-tuned modulation of the host cell transcriptome during *L. am* infection that relies on a pleotropic, yet highly regulated inhibition of inflammatory responses through antagonistic regulation of pro- and anti-inflammatory genes.

### Modulation of the level of acetylation of macrophage histone H3 during *L. am* infection

Since histone acetylation is crucial for the regulation of NF-κB-mediated inflammation (Ghizzoni et al., 2011), we next investigated if the dichotomic transcriptomic modulation of pro- and anti-inflammatory genes during *L. am* infection was linked to epigenetic regulation. Using an EpiTect ChIP qPCR Array, changes in histone H3 acetylation were assessed on a focused panel of 88 NF-κB associated gene promoters in two independent experiments (Figure S6 and Figure S6B). As expected from previous reports (Kapellos and Iqbal, 2016; Saeed et al., 2014), LPS treatment caused a strong global increase in H3 acetylation, known to be associated with transcriptional activation. This level of H3 acetylation was significantly reduced during *L. am* infection (Figure S6C). Even though infection alone did not affect the global H3 acetylation level in unstimulated macrophages (Figure S6C), parasite infection caused a reproducible hypoacetylation at the level of various pro-inflammatory promoters (Figure 4A), which correlated with the decreased abundance of the corresponding transcripts (see Figure 3), including TLR9, MYD88, or the NF-κB member RelA. Likewise, a correlation between H3 acetylation and transcript expression levels was observed among up-regulated genes, as for example for TNFAIP3, a NF-κB negative regulator playing a pivotal role in the termination of NF-κB-induced inflammation (Pujari et al., 2013). Thus, the dichotomy observed on transcript levels is largely reproduced on the level of H3 acetylation. Similar to the results obtained on transcript levels, such an epigenetic dichotomy was not observed in LPS activated macrophages for the promoters we analyzed: While the reduced acetylation observed for NF-κB positive regulators remained robust, no significant effect on inhibitory components was seen (Figure 4B), which explains the dampening rather than the abrogation of inflammasome activation. Nevertheless, *L. am* infection efficiently counteracted the increase in H3 acetylation caused by LPS, with H3 hypoacetylation demonstrated for example at the promoter level for the NF-κB inhibitor TNFAIP3. Overall, the global prevention of LPS-mediated H3 acetylation far exceeded the effect observed in unstimulated macrophages demonstrating the capacity of intracellular *L. am* to most efficiently interfere with the pro-inflammatory host cell response (Figure 4C).

**Figure 4:**
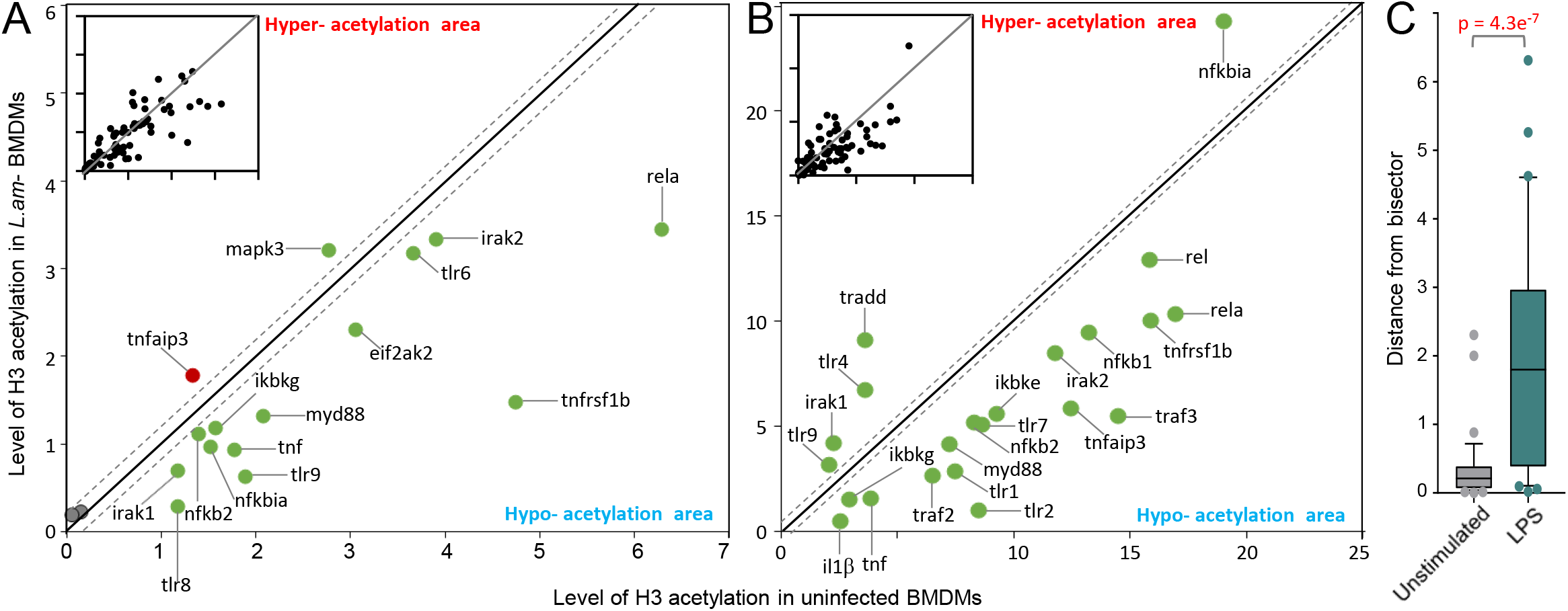
*L. am* infection modulates macrophage H3 acetylation at promoters of NF-κB-related genes. Chromatin ImmunoPrecipitation (ChIP) was performed on uninfected and *L. am*-infected BMDMs at 3 days PI, in presence or absence of LPS. Following nuclei isolation, chromatin shearing and immunoprecipitation with control and anti-acetylated histone H3 antibodies, qPCR assays were performed on chromatin preparations to measure enrichment of genomic DNA promoter sequence for genes associated with NF-κB signaling (EpiTect^®^ ChIP qPCR Array, mouse NF-κB Signaling Pathway). (A, B) Bi-parametric dot plots of the level of H3 acetylation in non-infected (X axis) and *L. am*-infected (Y axis) BMDMs in absence (A, mean values of n = 2 independent experiments) or presence (B, n = 1 experiment) of LPS stimulation. The inserts show the results obtained for all gene promotors analysed, while the main panels show the results for those gene for which abundance of their corresponding transcripts were analyzed previously by RT-qPCR (see Figure 3). The bisector is denoted by the diagonal black line. The unmodulated zone is comprised between the two dotted lines, that separates hypo- and hyper-acetylation zones. Green and red dots correspond to gene promoters associated to down- and up-modulated transcript abundance, respectively. (C) Box- and Whisker plots representing the distribution of the distance between H3 acetylation levels of unstimulated (A) and LPS-treated (B) in *L. am*-infected BMDMs and the corresponding bisector. Distributions have been calculated for genes whose transcript abundance has been assessed by RT-qPCR.

## Discussion

*Leishmania* has evolved molecular strategies to harness the phenotypic potential of its macrophage host cell, often with devastating consequences for infected individuals (Gollob et al., 2014; Kaye and Scott, 2011). Using primary murine macrophages infected with lesion-derived *L. amazonensis* amastigotes, we deliver here the first demonstration that this dominant human pathogen remodels the host cell chromatin during infection to establish permissive conditions for persistent, intracellular survival (Figure 5). We show that reduced H3 acetylation levels at host pro-inflammatory promoter genes correlates with decreased expression of crucial NF-κB and inflammasome activators. This finding reports a major new mechanism underlying the strong anti-inflammatory or immune subversive properties *Leishmania* exerts on its host cell irrespective of parasite species and tropism (de Freitas et al., 2016; Espitia et al., 2014; Kong et al., 2017; Liese et al., 2008; Soong, 2012; Stager et al., 2010) revealing a potential common epigenetic root for the immunopathologies underlying the different forms of clinical leishmaniasis (Gollob et al., 2014; Soong et al., 2012).

**Figure 5:**
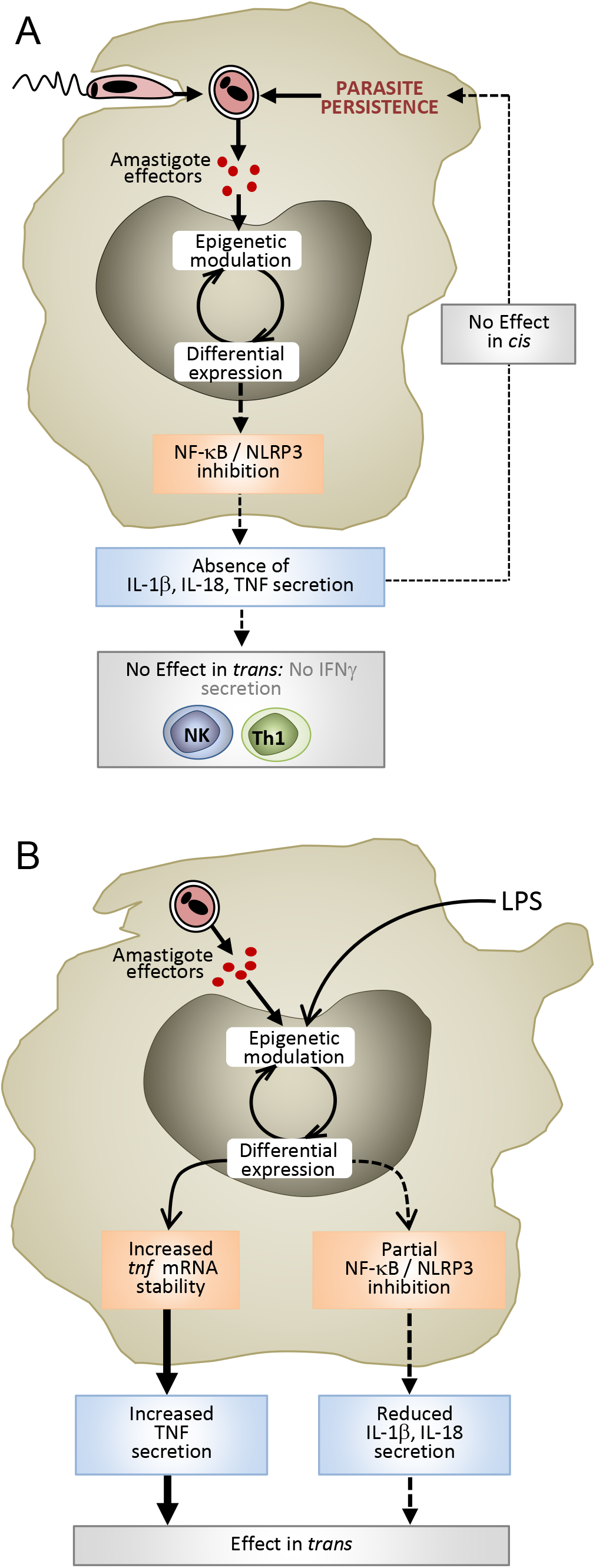
Models of the subversion of the NF-κB-NLRP3 immune axis induced by *L. am* amastigotes in BALB/c BMDMs. (A) Under steady-state conditions, unknown amastigote-released effectors modulate the host cell epigenetic profile, likely causing the important changes in transcript abundance we observed for the genes of the NF-κB and NLRP3 signaling pathways. Absence of pro-inflammatory IL-1β, IL-18 and TNF production allows for silent infection, undetected by non-infected bystander cells, thereby favoring the establishment of infection and early parasite survival. (B) In presence of bacterial products (i.e. LPS), the epigenetic effects caused by *Leishmania* amastigotes counteract those induced through TLRs and induce a paradoxical pattern of cytokine secretion, characterized by (i) increased TNF synthesis and secretion involving stabilization of the mRNA, and ii) decreased IL-1β and IL-18 production, which correlates with the transcriptional inhibition of positive regulators of the NF-κB and NLRP3 axes.

During microbial infections, the macrophage phenotype is regulated in a highly dynamic manner by a combination of cell-specific transcription factors and chromatin modifications, which adapt the cell’s transcriptional response to either promote or resolve inflammation (De Santa et al., 2009; Ehrt et al., 2001; Rolando et al., 2013). Various viral, prokaryotic and eukaryotic microorganisms have evolved strategies to interfere with host cell transcriptional and epigenetic regulation (Cock-Rada et al., 2012; Ding et al., 2010; Hamon et al., 2007; Han et al., 2012; Hari Dass and Vyas, 2014; Kinnaird et al., 2013; Lang et al., 2012; Leng et al., 2009; Marazzi et al., 2012; Marr et al., 2014; Rolando et al., 2013). Our data provide first evidence that *Leishmania* adopts a similar strategy causing a fine-tuned, dichotomic dysregulation of the macrophage inflammatory response, characterized by concurrent suppression of pro-inflammatory and upregulation of anti-inflammatory components of the NF-κB signaling and NLRP3 inflammasome pathways on transcriptional and epigenetic levels. Surprisingly, deactivating of these two major components of the host anti-microbial response (Bauernfeind and Hornung, 2013; Evavold and Kagan, 2018; Liang et al., 2004; Prochnicki et al., 2016; Rahman and McFadden, 2011; Smale, 2011) was stage specific and only observed for disease-causing amastigotes, an important aspect that escaped a recent investigation of inflammasome activation in response to insect-stage promastigotes (Lima-Junior et al., 2013).

While the underlying parasite effector mechanisms causing host cell immune suppression remain elusive, our study draws a detailed and complex picture of their impact on the host cell phenotype. We reveal a long-lasting (over three weeks) and global inhibition of the NF-κB-mediated pro-inflammatory response on the transcriptomics level, which extends previous findings on specific inhibition of this pathway early during infection involving cleavage of individual NF-κB members (Abu-Dayyeh et al., 2010; Calegari-Silva et al., 2009; Cameron et al., 2004; Gregory et al., 2008), the formation of Crel/P50 or P50/P50 dimers (Calegari-Silva et al., 2009; Guizani-Tabbane et al., 2004), the specific increase of key inhibitors of TLR-signaling such as TNFAIP3 or IRAKm (Srivastav et al., 2012; Srivastav et al., 2015), or the inhibition of TRAF3 degradation (Gupta et al., 2017). Thus, *L. amazonensis* not only avoids inflammasone activation early during infection allowing for establishment of macrophage infection, but reprograms its host cell into a safe haven permissive for long-term persistent infection and intracellular parasite growth. This phenotypic shift not only affects the host cells immune potential, but may also cause the profound metabolic effects observed in *Leishmania-infected* macrophages (Franca-Costa et al., 2015; Osorio y Fortea et al., 2009) that may feed-back on the macrophage phenotype and its polarization state (Jaramillo et al., 2011; Martin et al., 2012).

Our findings showcase *Leishmania* as a pivotal model system to probe macrophage plasticity. The previously described *Leishmania*-induced metabolic re-tooling of the host cell directly supports the anti-inflammatory effect of parasite infection, likely through establishing a M2-like macrophage phenotype - a polarization profile known to be involved in the resolution of inflammation and characterized by up-regulation of inhibitors of the NF-κB signaling pathway and the decreased expression of IL-1β, IL-18 and inflammasome components (Awad et al., 2017; Osorio y Fortea et al., 2009). Understanding macrophage plasticity using intracellular pathogens such as *Leishmania* and ultimately harnessing this process by pharmacological or immunological intervention opens exciting new venues for therapy. Our novel insights into *Leishmania* epigenetic, anti-inflammatory control can stimulate the development of new therapeutic options, for example mimicking the parasite’s suppressive effect on the NF-κB pathway, which is a major focus of curative intervention given its role in chronic inflammatory diseases (Durand and Baldwin, 2017; Herrington et al., 2016; Lin et al., 2017; Zeligs et al., 2016). Likewise, our findings open exciting new venues for host-directed, anti-leishmanial intervention strategies targeting the host cell epigenome, which has been recently proposed as a promising new way to guard against the selection of drug resistant parasites (Kumar et al., 2017; Lamotte et al., 2017; Prieto Barja et al., 2017). Reversing the parasite-driven epigenetic effect may rescue the host cell’s antimicrobial potential and establish a non-permissive, anti-leishmanial macrophage phenotype, or restore normal macrophage metabolic functions incompatible with intracellular parasite proliferation (De Muylder et al., 2016; Mukherjee et al., 2014; Osorio y Fortea et al., 2009).

In conclusion, we provide first evidence for remodeling of the macrophage chromatin during *Leishmania* infection that impacts on the host cells immune effector functions and metabolism establishing permissive conditions for intracellular parasite survival. Whether the *Leishmania*-induced changes in the host cell epigenome are cause or consequence of the observed phenotypic response remains to be elucidated. However, the correlation we observed between histone H3 hypoacetylation and transcriptional downregulation for pro-inflammatory surface receptors (TLR4, TNFR), signaling molecules (MYD88, RIPK2), transcription factors (RelA, RelB, NF-κB1, NF-κB2) and cytokines (IL-1β, TNF) suggests that chromatin remodeling is the cause of the macrophage phenotypic change. Future studies investigating macrophage/ *Leishmania* interaction at systems level combining transcriptomics, epigenetics and metabolomics analyses will solve this important open question and uncover interesting new candidates for host-directed, anti-leishmanial therapies.

## AUTHOR CONTRIBUTIONS

H.L., E.P., T.R., K.K., P.N., and Y.X. performed research. and analyzed data. H.L., E.P., T.R., K.K., P.N., Y.X., N.A, H.V. and G.B. analyzed data. R.W. gave personal reagents. H.L., E.P., and G.F.S. designed research and wrote the paper. All authors read and approved the final manuscript.

## ACKNOWLEDGEMENTS

This project was supported by a fund of the Institut Pasteur International Direction to the International Mixed Unit ‘Inflammation and *Leishmania* infection’. We would like to thank Drs Genevieve Milon and Jean-Marc Cavaillon for support and helpful discussions, and Drs Emmanuel Laplantine and Anastassia V. Komarova for providing antibodies. The UtechS PBI/C2RT is part of the FranceBioImaging infrastructure supported by the French National Research Agency (ANR-10-INSB-04-01, Investments for the Future) and is supported by the Conseil de la Region Ile-de-France (program Sesame 2007, project Imagopole, S. Shorte) and the Fondation Française pour la Recherche Médicale (Programme Grands Equipements [N.A.])

## DECLARATION OF INTEREST

The authors declare no competing financial interests.

## SUPPLEMENTAL FIGURE LEGENDS

**Figure S1, related to figure 1:**
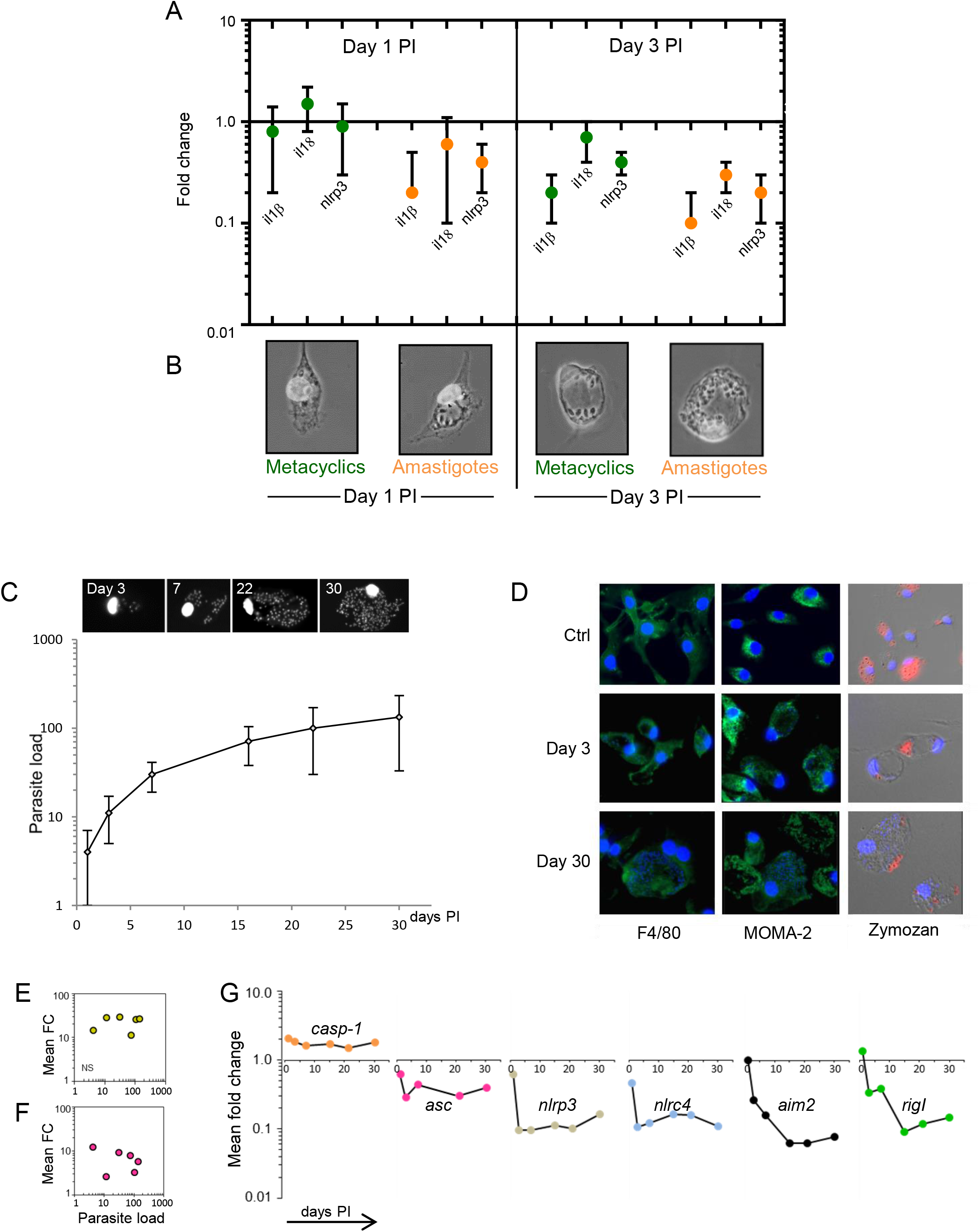
Monitoring macrophage subversions in short- and long-term cultures. The delayed transcriptional subversion in macrophages infected by *L. am* metacyclic promastigotes vs amastigotes was analysed in parts A and B. BMDMs were infected with *L. am* metacyclic promastigotes (metacyclics) or lesion-derived amastigotes and cultured for one and three days. **(A)** Transcript modulations for *il1β, il18* and *nlrp3* were assessed by RT-qPCR (mean fold changes ± SEM) for metacyclic promastigotes- (green symbols) and amastigotes- (orange symbols) infected cultures (n=3) for three. **(B)** Pictures of representative infected BMDMs. Note the delay of parasitophorous vacuole formation in the macrophages infected with metacyclic promastigotes compared to amastigote-infected cells. The phenotypic and transcriptional characterization of long-term cultured BMDMs infected with *L. am* amastigotes was analysed (parts C-G). BMDMs were infected with *L. am* amastigotes and cultured up to 30 days PI. (C) Monitoring of parasite load in infected macrophages. Representative images of infected BMDMs stained with Hoechst 33342 are shown in the upper panel (epifluorescence microscopy). Parasite load corresponds to the number of Hoechst-stained parasite nuclei per cell, determined after counting a minimum of 100 macrophages (Means ± SEM, one representative experiment). (D) Immuno-phenotyping and evaluation of the phagocytic capacity of infected BMDMs. Specific macrophage lineage markers F4/80 and MOMA-2 (green staining) were visualized at day 3 (Ctrl and *L.am* infected BMDMs) and 30 PI by epifluorescence microscopy. Phagocytosis ability was assessed with Texas Red-conjugated zymosan particles (red staining) at the same time points. Phase contrast and immunofluorescence image superposition shows healthy macrophages with large PVs and numerous intravacuolar amastigotes. BMDM and parasite nuclei were identified after DNA staining with Hoechst 33342 (blue staining). (E, F, G) Kinetics analysis of inflammatory cytokines and inflammasome-related transcripts. RT-qPCR analyses were performed at 1, 3, 7, 16, 22 and 30 days PI. Data are represented as expression fold changes for *il1α* (E), *tnf* (F) and various inflammasome-related transcripts (G). No correlation was observed between the parasite load and the fold changes for *il1α* and *tnf*

**Figure S2, related to figure 1:**
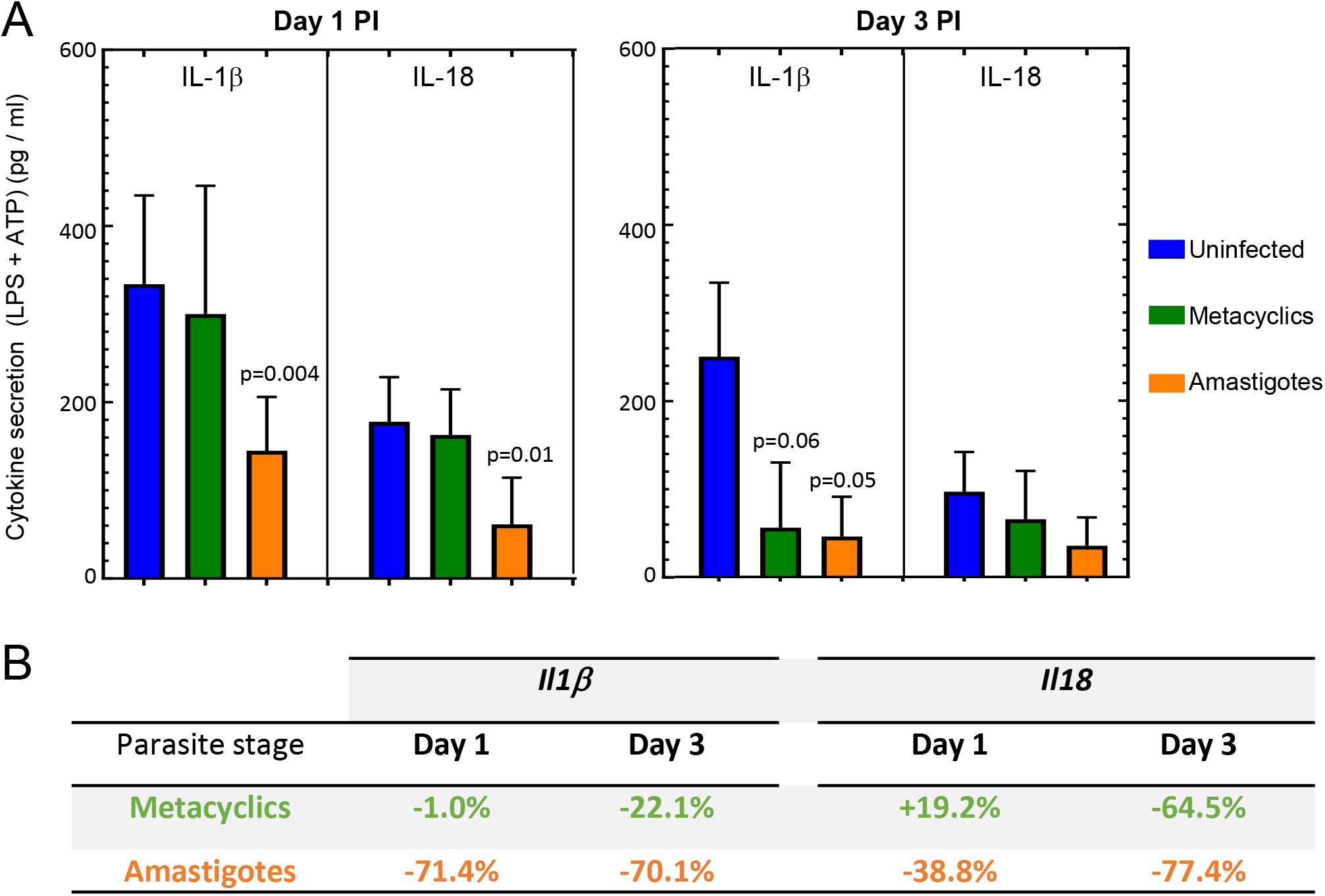
*L. am* amastigotes induce specific transcriptional subversions in LPS/ATP stimulated macrophages. BMDMs were infected with *L. am* purified metacyclic promastigotes (metacyclics) or amastigotes and cultured for one and three days before sequential stimulation with LPS (4 hours) and ATP (2 hours). Cytokines were quantitated in the supernatant and the corresponding transcripts were analysed in the corresponding cell samples. Results were obtained on n = 3 independent experiments. **(A)** Quantitation of IL-1β and IL-18 secretion by ELISA. Mean ± SEM values are displayed in blue (uninfected control), green (metacyclic promastigotes) and orange (amastigotes) histograms. **(B)** Analysis of *il1β* and *il18* transcripts in the corresponding samples by RT-qPCR. Modulations of *il1β* and *il18* transcript abundance in response to infection by metacyclics or amastigotes are displayed by the percentage of modulation versus non-infected cultures.

**Figure S3, related to Figure 1:**
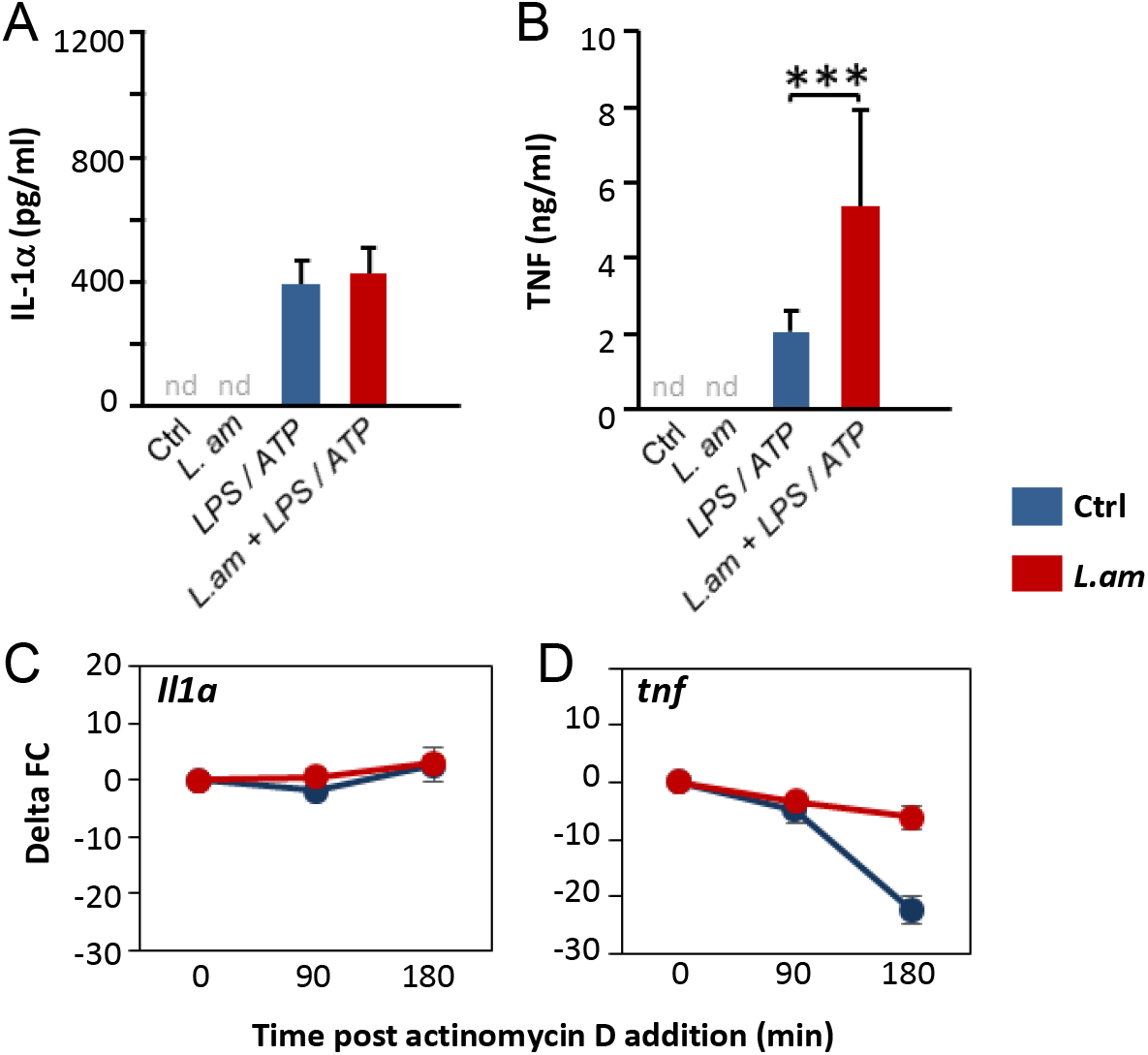
Modulation of IL-1α and TNF secretion in LPS / ATP stimulated BMDMs after *L. am* infection. BMDMs were infected or not by *L. am* amastigotes for 3 days and sequentially stimulated with LPS (4 hours) and ATP (2 hours). (A, B) Quantitation of IL-1α **(A)** and TNF **(B)** in supernatants by ELISA. Blue and red bars represent cytokine levels for uninfected and infected BMDMs respectively (mean +/− SEM, n = 3 - 6; nd, not detected). Significant p value: p < 0.001 (***). **(C, D)** Analysis of transcript stability after 4 hours of LPS stimulation and further addition of the transcription inhibitor actinomycin D for 90 and 180 min. Transcripts stability was analyzed for *il1 α* **(C)** and *tnf* **(D)** in uninfected (blue line) and infected (red line) BMDMs. Data are represented as differences in fold changes (Delta FC) between transcript abundance of untreated BMDMs and actinomycin D-treated BMDMs. *tnf* transcript stabilization is reflected by the stability of the delta FC in presence of *Leishmania* amastigotes (red line).

**Figure S4, related to Figure 2:**
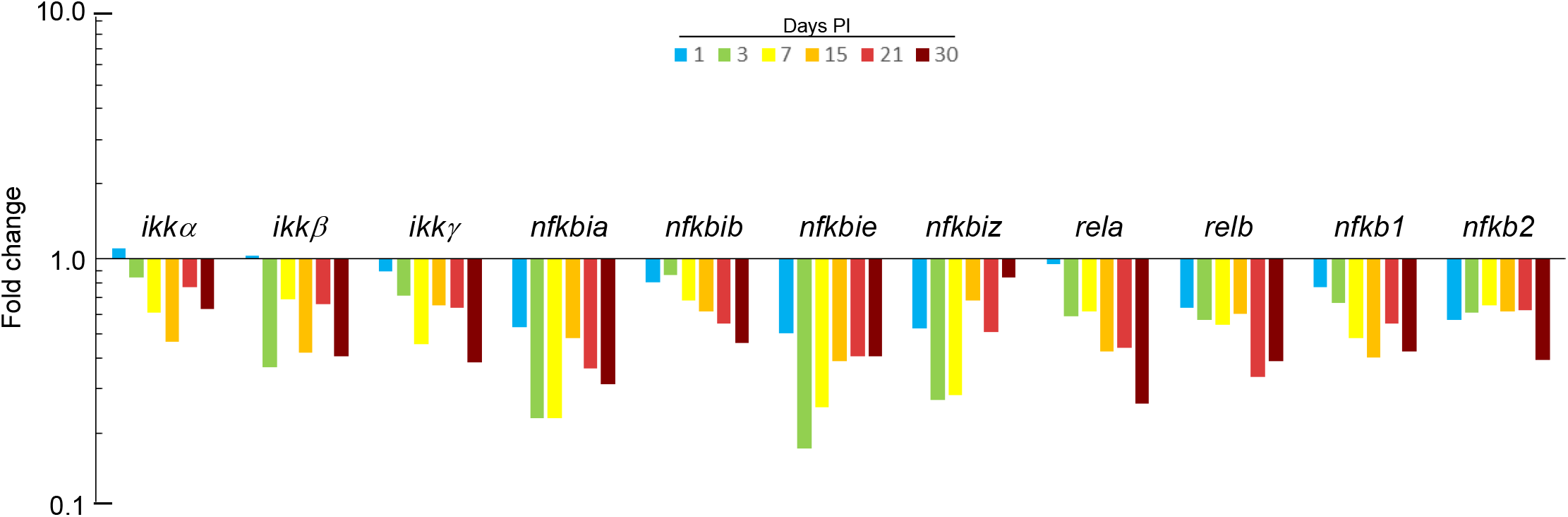
Long-term transcriptional subversion of NF-κB signaling members induced by *L. am* amastigotes. BMDMs were infected with *L. am* amastigotes and transcriptional profiling of NF-κB-related genes in *L. am*-infected and uninfected BMDMs was performed by RT-qPCR at different time points PI. Fold changes using uninfected BMDMs as calibrator are shown for one representative experiment (n= 2).

**Figure S5, related to Figure 3:**
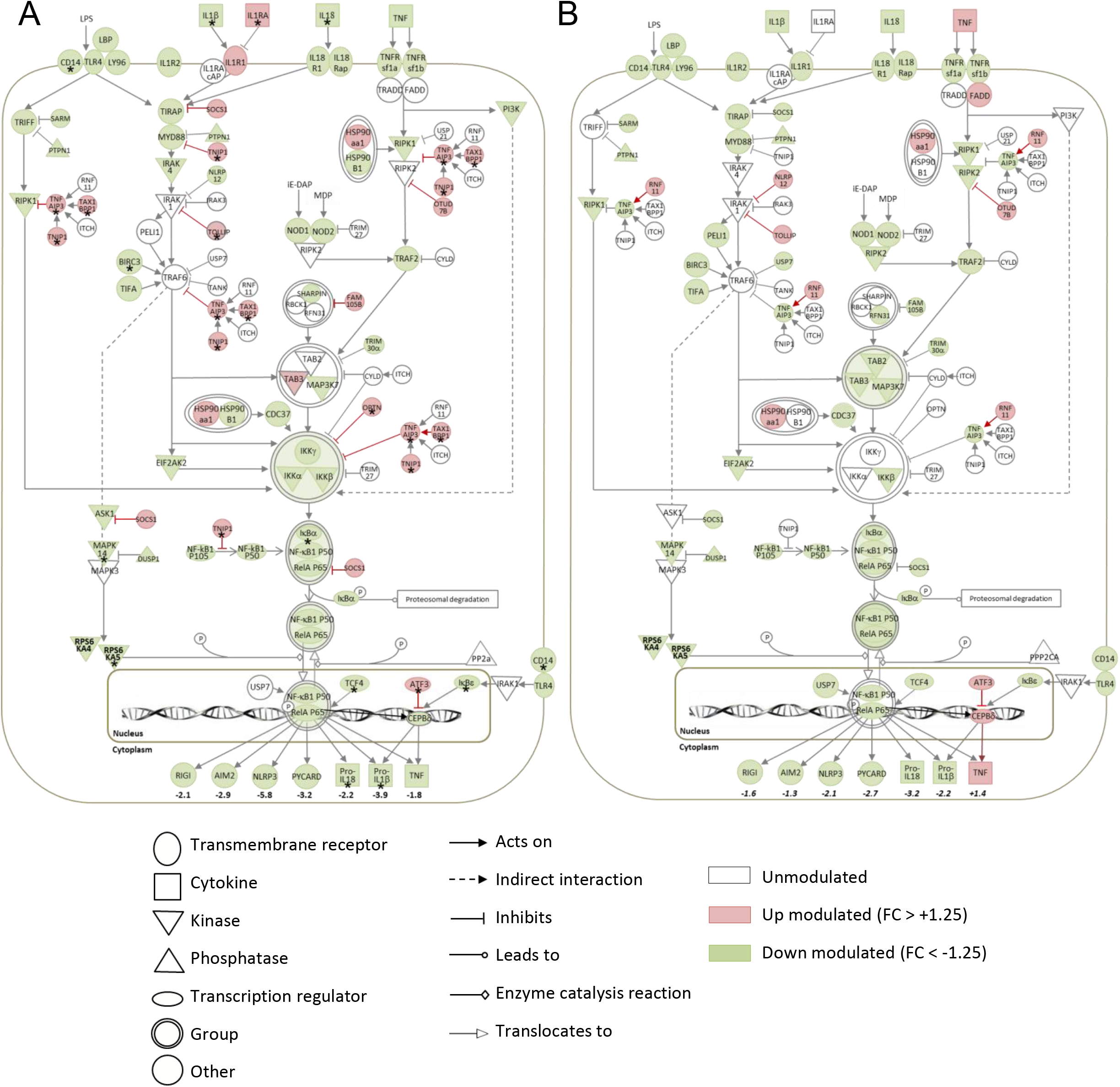
Schematic view of the transcriptional profiling of the modulations of NF-κB-NLRP3 signaling pathway in infected BMDMs in absence (A) or presence (B) of LPS and ATP. BMDMs were infected or not by *L. am* amastigotes for 3 days. Transcriptional profiling was performed by RT-qPCR. Fold changes calculated between infected and uninfected BMDMs are represented by the color code with red for up regulated transcripts (FC > +1.25) and green for down-regulated transcripts (FC<-1.25) (n = 3 - 8 independent experiments). The asterisks marks transcripts that were significantly modulated in our previous study at day one PI as evidenced using the Affymetrix technology (Osorio y Fortéa et al., 2009). Figures correspond to FC values for inflammasome-related transcripts.

**Figure S6, related to figure 4:**
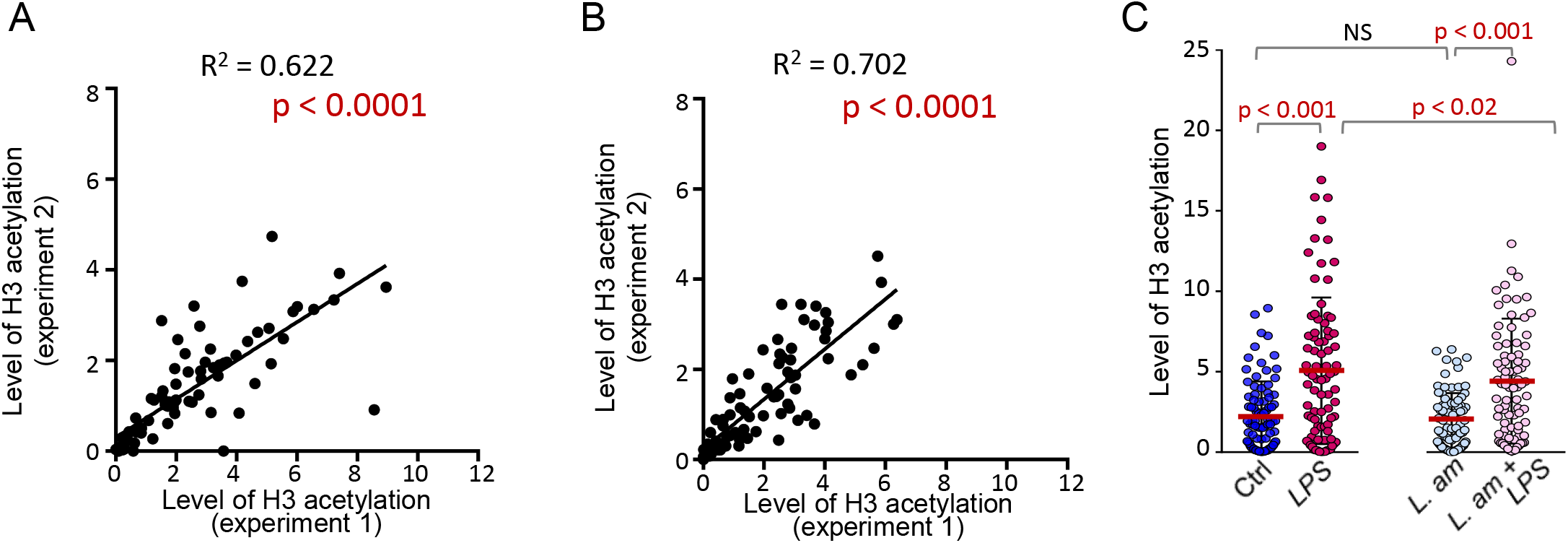
Analysis of the level of H3 acetylation in BMDM samples.. BMDMs were infected or not with *L. am* amastigotes for 3 days and stimulated for 4 hours with LPS. Following nuclei isolation, chromatin shearing and immunoprecipitation with anti-acetylated histone H3 antibodies, qPCR assays were performed to measure enrichment of genomic DNA promoter sequences for the genes of the NF-κB pathway (EpiTect® ChIP qPCR Array, mouse NF-κB Signaling Pathway). H3 acetylation level data for every gene were expressed as percentage of input in uninfected **(A)** and *L. am*-infected **(B)** BMDMs and displayed for 2 independent experiments. Coefficient of determination and p values from linear regression are shown. **(C)** Histone H3 acetylation levels expressed as percentage of input corresponding to the sample before immunoprecipitation for ctrl, LPS-treated, *L.am-* infected and *L.am-* infected treated with LPS samples. The dot plot shows the mean percentage of input (n = 2 independent experiments). Statistical analysis was performed using the Wilcoxon matched-pairs and signed rank test.

## STAR★METHODS

### EXPERIMENTAL MODEL AND SUBJECT DETAIL

#### Ethics statement

Six-week-old female BALB/cJRj and Swiss *nu/nu* mice were purchased from Janvier (Saint Germain-sur-l’Arbresle, France) or from Shanghai Laboratory Animal Center (SLAC). All animals were housed in A3 animal facilities according to the guidelines of Institut Pasteur and the “Comité d’Ethique pour l’Expérimentation Animale” (CEEA) and protocols were approved by the “Ministère de l’Enseignement Supérieur; Direction Générale pour la Recherche et l’Innovation” under number 2013-0047 and by the Animal Care and Use Committee at Institut Pasteur of Shanghai Animal Care.

#### Isolation and culture of bone marrow-derived macrophages

Bone marrow cell suspensions were recovered from tibias and femurs of BALB/c mice in DMEM medium (Gibco, Life technologies) and cultured in medium complemented with mouse recombinant colony stimulating factor 1 (mrCSF-1, ImmunoTools) (de La Llave et al., 2011). One million cells per ml were incubated in bacteriologic petri dish (Corning Life Science) at 37°C in a 7.5% CO2 atmosphere for 6 days with 75 ng/mL mrCSF-1. After detachment with 25 mM EDTA, 2 million macrophages were seeded per well in 12-well plates in 2 ml culture medium containing 20 ng/mL mrCSF-1. Inflammasome activity was analyzed after sequential treatment with 500 ng/ml LPS for 4 hours and 5 mM ATP for 2 hours. Proteasomal protein degradation was studied in presence of 10 μM carbobenzoxy-Leu-Leu-Leucinal (MG132) during 4 hours.

#### Parasite isolation and BMDM infection

*mCherry* transgenic, tissue-derived amastigotes of *Leishmania amazonensis* strain LV79 (WHO reference number MPRO/BR/72/M1841) were isolated from infected footpads of Swiss nude mice (Lecoeur et al., 2010). Amastigotes were added at a ratio of 4 amastigotes per macrophage, reaching 95% of infection (fluorescence microscopy analysis of Hoechst-stained samples). Metacyclic promastigotes were isolated from stationary phase amastigote-derived promastigote cultures on a discontinuous Ficoll gradient (Spath and Beverley, 2001) and added to BMDMs at a multiplicity of infection (MOI) of 8:1. Cells were cultured at 34°C for 3 to 30 days post-infection (PI). The supernatant was replaced once a week by fresh complete medium containing 20 ng/ml mrCSF-1. BMDMs were allowed to ingest heat-killed, tissue-derived amastigotes obtained after 20 min treatment at 45°C, or inert latex beads (5 μm particle size, Sigma-Aldrich) at a bead to macrophage ratio of 20:1. Phagocytic activity was determined using Texas red-labeled zymosan (ThermoFisher Scientific).

### METHOD DETAILS

#### Western blotting

BMDMs were lysed in RIPA buffer (R0278, SIGMA) containing anti-proteases and anti-phosphatases inhibitors (MS-SAFE, SIGMA). Proteins were resolved by SDS–PAGE on 4–12% Bis-Tris NuPAGE gels in MOPS buffer and electroblotted onto polyvinylidene difluoride membranes. Membranes were blocked with 5% fat-free milk in Tris-buffered saline containing 0.25% Tween 20 and then probed overnight at 4°C with anti-NLRP3 (MAB7578, R&D Systems), NLRC4 (NB100-56142, Novus Biologicals), AIM2 (ab93015, abcam), RIG-I (3743, D14G6 clone, Cell Signaling), caspase-1 p20 (AG-20B-0042, Adipogen Life Sciences), IκBα (ab32518, abcam), IκBβ (AM8109a, 62AT216 clone, Abgent), phospho IκBα (MA5-14857, clone J10.3,ThermoFischer scientific), IKKγ (sc-8330, Santa Cruz Biotechnology) and β-actin (4970, Cell Signaling) antibodies.

To detect the ASC pyroptosome, BMDMs were lysed in 50 mM Tris pH 7.5, 150 mM NaCl, 1% NP-40, supplemented with MS-SAFE, and sheared 10 times through a 21-gauge needle. Lysates were cross-linked with 4 mM disuccinimidyl suberate and resolved by electrophoresis on 12% SDS-PAGE. Immunoblotting was performed with anti-ASC antibody (sc-22514-R, Santa Cruz Biotechnology).

To detect released IL-1β and caspase-1, soluble proteins from cell supernatants were precipitated with methanol and chloroform, separated by SDS-PAGE, transferred to nitrocellulose membranes and immunoblotted with anti-IL-1β (sc-7884; Santa Cruz Biotechnology) or caspase-1 p20 (AG-20B-0042, Adipogen) antibodies.

Following incubation with peroxydase-conjugated secondary antibodies, membranes were revealed by SuperSignal West Pico reagent (ThermoFisher Scientific) in a high resolution PXi machine (Syngene). Relative protein expression was calculated by densitometric analysis using the ImageJ software. Ratios between integrated density values obtained for the target protein and β-actin were calculated. Fold changes were expressed using the control sample as a calibrator, with control values of uninfected and unstimulated samples being set to 1.

#### RNA extraction and transcriptional analyses by real time quantitative PCR (RT-qPCR)

Total RNA isolation, quality control and reverse transcription were performed as previously described (de La Llave et al., 2011). RT-qPCR was carried out in 384-well PCR plates (Framestar 480/384, 4titude, Dominique Dutscher) using the iTaq™ Universal SYBR^®^ Green Supermix (Bio-Rad) and 0.5 μM primers with a LightCycler^®^ 480 system (Roche Diagnostics, Meylan, France). Primer information for every target tested by qPCR is detailed (Table S1-3). Crossing Point values (Cp) were determined by the second derivative maximum method of the LightCycler^®^ 480 Basic Software. The relative expression software tool (*REST©-MCS*) (Pfaffl et al., 2002) was used to determine relative expression as fold change (FC) values and normalization was performed using the geometric mean of *ywhaz* and *rpl19* quantities as determined by using GeNorm and Normfinder programs following the expression analysis of various candidate control genes (de La Llave et al., 2011). For statistical analysis of gene expression levels Cp values were first transformed into relative quantities (RQ). Then, normalized RQ (NRQ) were calculated by dividing the RQ of each gene of interest by the normalization factor i.e. the geometric mean of RQ of *ywhaz* and *rpl19, the best combination of* reference genes in our experimental conditions (de La Llave et al., 2011). Nonparametric Kruskal-Wallis tests were performed on Log transformed Normalized Relative Quantity values.

The modulation of macrophage RNA stability by *L. am* amastigotes was analyzed after blocking or not the transcription with 5 μg / ml Actinomycin D (AD, A9415, Sigma) in uninfected and infected samples. RNA were analyzed after 90 and 180 minutes of AD treatment following a 4 hr LPS stimulation period. The modulation of RNA stability by *L. am* amastigotes was determined by comparing the FC values obtained between AD-treated versus AD-non treated cells samples.

Heatmaps of the Fold-Changes on the log2 scale (RT-qPCR) and evaluation of the differences in H3 acetylation between infected and uninfected samples were performed using R (version 3.4.3.) and home-made scripts based on native graphical functions.

#### Cytokine quantitation in culture supernatants

Cytokines were quantified in the supernatant using mouse instant ELISA kits (for IL-1β and TNF, eBioscience) or classical mouse ELISA kits for IL-1α (eBioscience) or IL-18 (R&D system Europe).

#### Microscopic and immunofluorescence analyses

BMDMs were seeded in complete medium containing 20 ng/ml mrCSF-1 either on glass slides at a density of 1.5×10^5^ cells (24-well plates) or 5×10^4^ cells (96-well plates) per well. BMDMs were fixed and treated as previously described (Prina et al., 2004) and incubated in 0.1% saponin buffer containing 0.25% gelatin and anti-F4/80 (MCA497 ABDSEROTEC), MOMA-2 (MCA519 AbD Serotec) and RelA polyclonal antibodies (ab16502, abcam). Stainings were revealed using appropriate secondary antibodies and nuclear staining was peformed with Hoechst 33342 (ThermoFisher Scientific). Image acquisition and analysis for glass coverslips were performed using the upright ZEISS Axio Imager 2 and the Zen Imaging Software. Image acquisition for 96-well plates was performed on the automated OPERA QEHS spinning disk confocal microscope (Perkin Elmer Technologies). Analysis routines of the Columbus software package (PerkinElmer Technologies) were used for automated scoring of fluorescence signals for (i) nuclear localization of p65 (RelA), which was delineated by nuclear counter stain with Hoechst 33342, and (ii) parasite detection using the mCherry signal. Twenty-eight fields per well were acquired allowing at least 800 cells to be analyzed per condition.

#### Chromatin isolation, Chip Analysis and qPCR

BMDMs were seeded in 100 mm tissue culture dishes. At day 3 PI, chromatin preparation, isolation and shearing from BMDMs were performed using a modification of a described procedure (Arrigoni et al., 2016). BMDMs were washed with pre-warmed (34°C) protein-free medium, fixed with 1% methanol-free formaldehyde at 34°C for 10 min, incubated 10 min in 125 mM Glycine at 4°C. Cells were scraped into Nuclei EXtraction by SONication (NEXSON) buffer (5 mM PIPES pH 8, 85 mM KCl, 0.5% Igepal CA-630) containing 2 x Halt protease inhibitor cocktail (Thermo Scientific). Samples were sonicated in 1.5 ml Eppendorf tubes using the Bioruptor device (Diagenode) at “low power” with two cycles (30” on 30” off), and the macrophage nuclei isolation was microscopically evaluated before collection of the nuclei by centrifugation for 5 min at 4°C at 2000g and resuspension in shearing buffer (10 mM Tris-HCl pH 8, 0.1% SDS, 1 mM EDTA, 2 x Halt protease inhibitor cocktail). Chromatin was sheared by three 5 min ultra-sound pulses at 4°C with sonicator set at “high power” to obtain 100-500 bp size fragments. Chromatin immunoprecipitations were performed with the EpiTect^®^ ChIP OneDay Kit (Qiagen) using control IgGs and antibodies against acetylated histone H3 (H3ac) (EpiTect^®^ ChIP Antibody Kit). The EpiTect^®^ ChIP qPCR Array Mouse NF-κB Signaling Pathway was applied on (i) 1/100 of the input material before immunoprecipitation as positive control, (ii) negative control IgGs from normal non-immune serum for background assessment, and iii) H3ac fractions to determine epigenetic status for genes of the NF-κB pathway. RT-qPCR data were obtained on a LightCycler^®^ 480 system (Roche Diagnostics, Meylan, France) using the 384-well template (GM-025G, Qiagen), and were expressed as percent of input for every target gene in uninfected and infected BMDMs. The distance between H3 acetylation levels in NF-κB related promoters and the corresponding bisector was analyzed for *L. am*-infected versus non-infected BMDMs, in presence or not of LPS.

### QUANTIFICATION AND STATISTICAL ANALYSIS

#### Statistical analysis

Statistical analysis of gene expression levels determined by RT-qPCR were performed in SigmaPlot Software (SigmaPlot for Windows Version 11.0, Build 11.2.0.5). Other statistical analyses were performed by the nonparametric Wilcoxon rank-sum test using the GraphPad Prism 7.03 software.

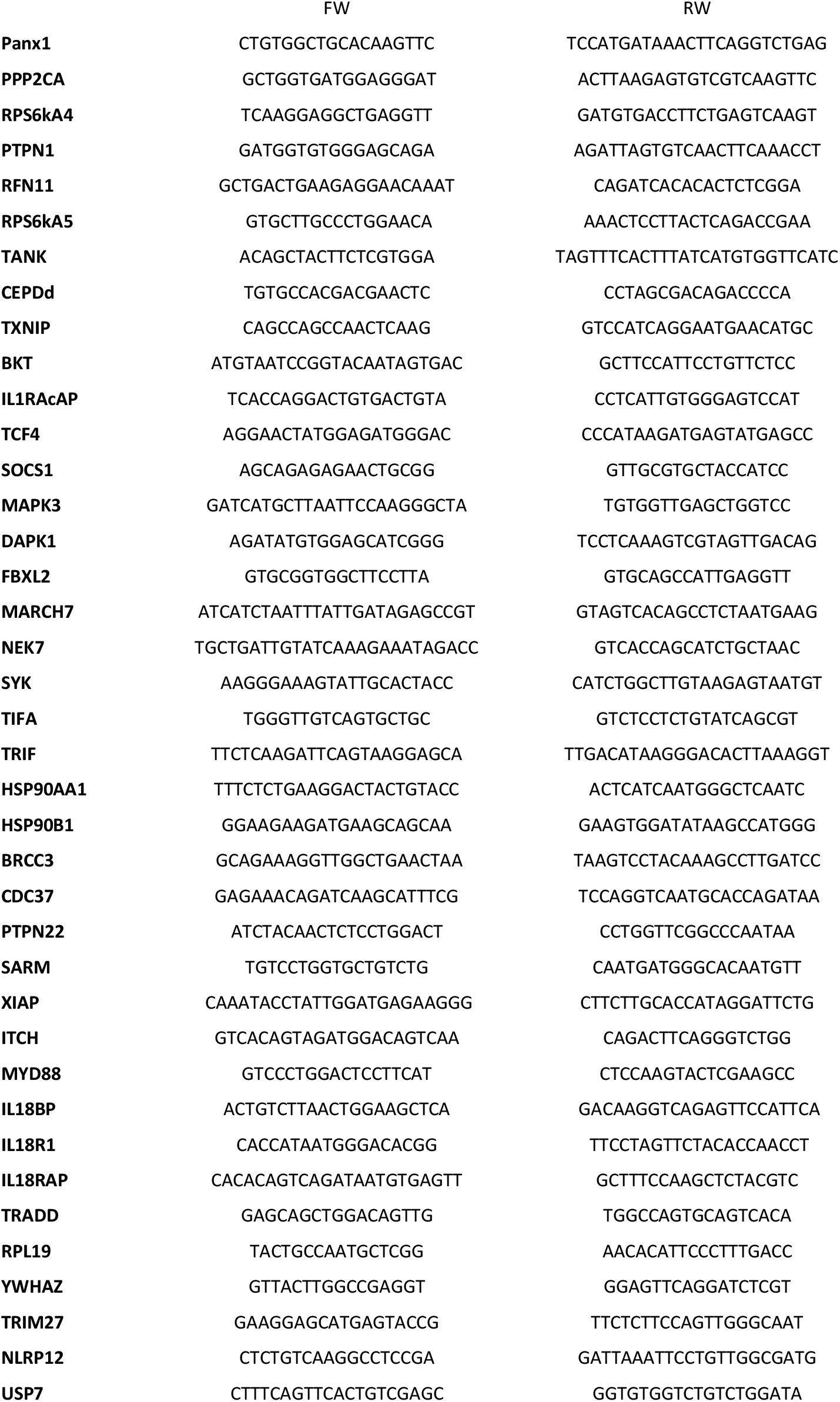
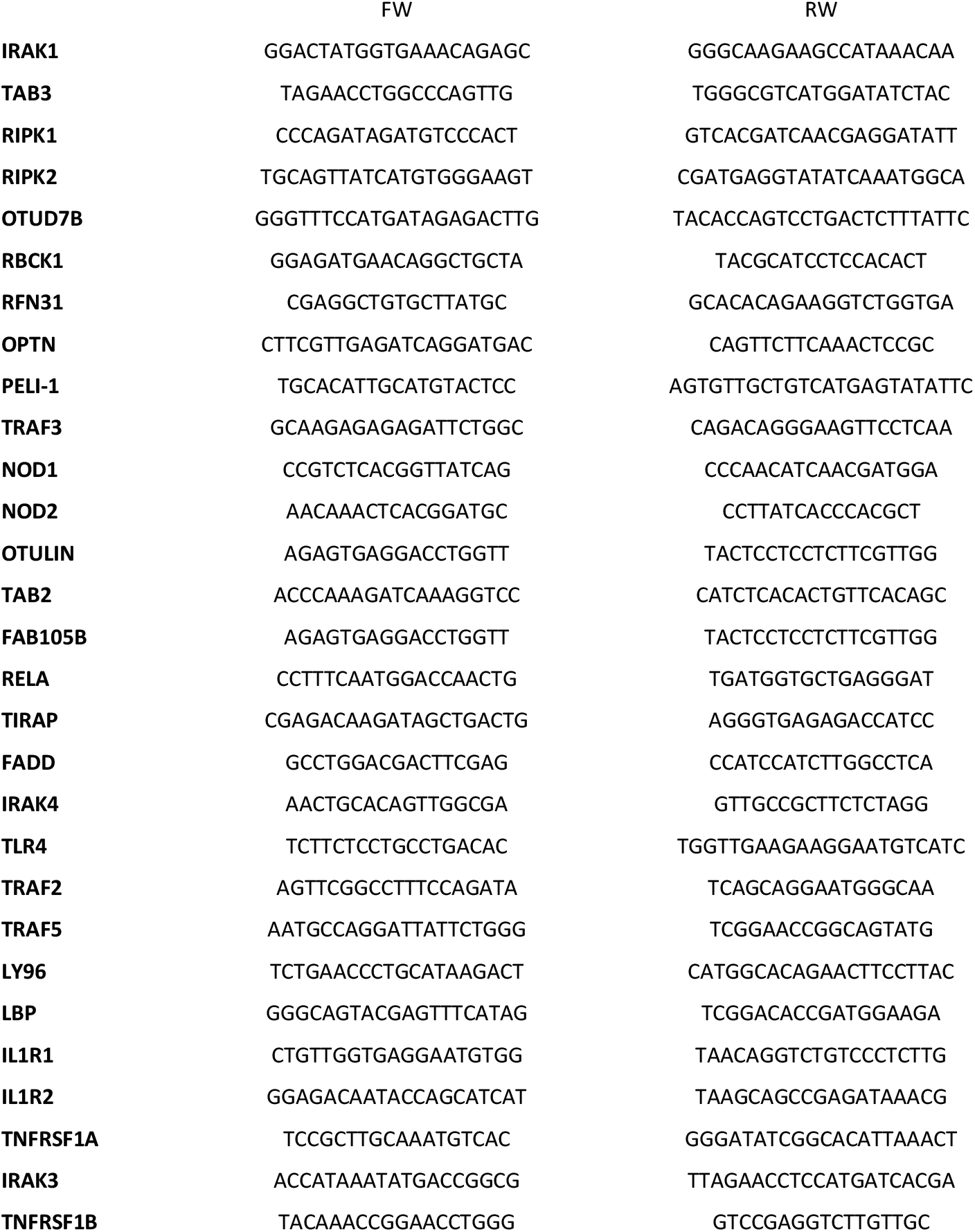
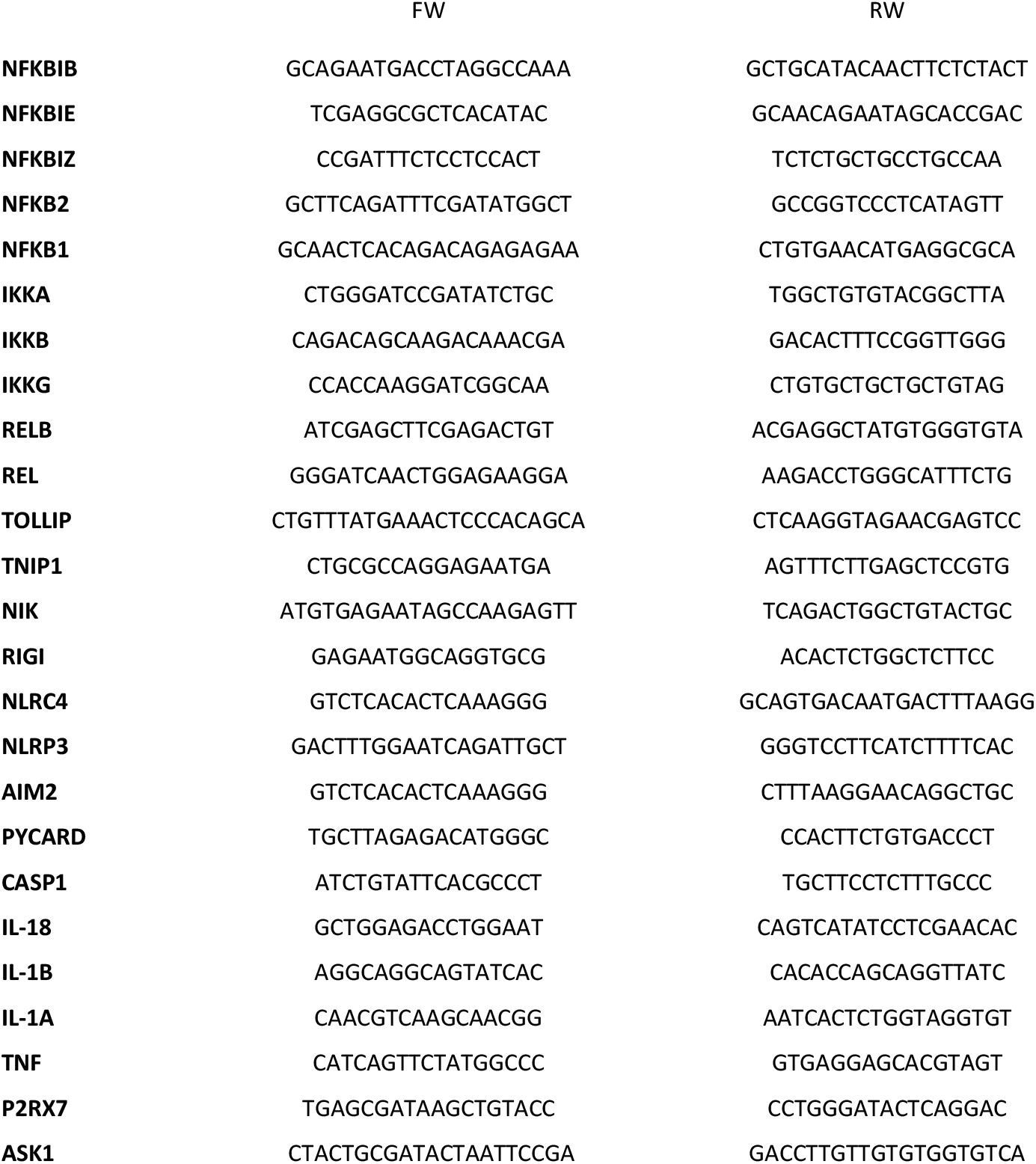

